# Benzo(a)pyrene regulates chaperone-mediated autophagy via heat shock protein 90

**DOI:** 10.1101/2022.06.08.495280

**Authors:** Min Su, Shuhong Zhou, Jun Li, Nan Lin, Tao Chi, Mengdi Zhang, Xiaoli Lv, Yuxia Hu, Tuya Bai, Fuhou Chang

**Affiliations:** School of Pharmacy, Inner Mongolia Medical University, Hohhot, China; School of Basic Medicine, Inner Mongolia Medical University, Hohhot, China; GLP Center of Inner Mongolia Medical University, Hohhot, China; Inner Mongolia New Drug Screening Engineering Research Center, Hohhot, China

**Keywords:** Benzo(a)pyrene, Chaperone-mediated autophagy, Heat shock protein 90, Heat shock cognate 70, Lysosomal-associated membrane protein type 2 receptor

## Abstract

The Benzo(a)pyrene (Bap) exposure induced oxidative damage, DNA damage and autophagy. To explore the molecular mechanism of BaP-induced autophagy. In these studies, we first found that heat shock protein 90 (*HSP90*), heat shock cognate 70 (*HSC70*) and lysosomal-associated membrane protein type 2 receptor (*Lamp-2a*) expressions of C57BL mice lung tissue and A549 cells exposed to BaP were significant increase, as well as Bap induced DNA double-strand breaks (DSBs) and activated DNA damage responses, as evidenced by *comet* assay and γ-H2AX foci analysis in A549 cells. Our results demonstrated BaP induced CMA and caused DNA damage. Next, we knocked down HSP90 expression by the HSP90 Inhibitor, NVP-AUY 922, exposed or HSP90α shRNA lentivirus transduction in A549 cells. HSC70 and Lamp-2a expressions of these cells exposed to BaP were not significant increase, which showed that BaP inducted CMA was mediated by HSP90. Further, HSP90α shRNA prevented BaP induced of Bap which suggested BaP regulated CMA and caused DNA damage by HSP90. Our results elucidated a new mechanism of BaP regulated CMA through HSP90.

## 1. Introduction

Autophagy is a cellular degradation and recycling process that is highly conserved in all eukaryotes. Three forms of autophagy have been described: macroautophagy, microautophagy and chaperone-mediated autophagy (CMA) in mammalian cells [1]. Selectivity lies in the chaperone heat shock cognate 71 kDa protein (HSC70) recognizing a pentapeptide motif (KFERQ-like motif) in the protein sequence, and form the chaperone/substrate complex [2]. Subsequently, heat shock protein 90 (HSP90) recognizes the complex and prevent formation of lysosomes[3,4].

Aryl hydrocarbon receptor (AhR) is complexed with HSP90 and p23 in the cytoplasmic. Benzo(a)pyrene (Bap) is a ligand of AhR, the prototypical polycyclic aromatic hydrocarbon carcinogen. Upon Bap binding, HSP 90 and AhR separates, and HSP90 is released in the cytoplasm. A study reported that smoking could cause increased expression of autophagy-related (ATG) genes [5,6]. It is well known one product of the tobacco products of Benzo(a)pyrene (Bap), so our study explore whether Bap affects autophagy and HSP90 is involved in the regulation of CMA caused by Bap.

To date, there have been a multitude of studies that the Bap partake in reactions leading to oxidative damage. And Bap metabolite of Benzo(a)pyrene diol epoxide (BPDE) and DNA form adducts. Oxidative damage and adducts lead to DNA double-strand break (DSB), causing gene instability induced autophagy [7,8]. Whether HSP90 participated in the CMA caused by DNA double-strand break, is our concern another focus.

Based on a multitude of reviews and studies, we propose two hypotheses that BaP binding to AhR, HSP90 is dissociate from AhR, and is involved in the recognition of substrates by HSC70, thereby affecting autophagy. Bap leads to gene instability and inducing autophagy by causing oxidative damage and forming the adducts of BPDE and DNA. HSP90 also plays a regulatory role in this process, that is, HSP90 is involved in the regulation of DNA damage induced by Bap and thus promotes the process of CMA.

## 2. Materials and methods

### 2.1 Animals model

C57BL mice weighing 25–30g (Institute of Animals of Inner Mongolia University) were housed in a specific pathogen-free environment. Mice in the Bap group were fed Bap (Sigma, City of Saint Louis, USA), solubilized in corn oil at a dose of 25.3mg/kg, twice per week, while the control group was given an equal volume of corn oil [9]. All procedures were approved by the Institutional Animal Care and Use Committee (IACUC) of Inner Mongolia Medical University.

### 2.2 MTT method

The density of 5 × 10^4^ cells/well of A549 cells were inoculated on 96-well plates. Then, the cells were treated with different concentrations (0.00µM, 0.01µM, 0.1µM, 1µM, 10µM) BaP for 24h, 48h, and 72h. MTT reaction mix (20µL of 5mg/mL) were added to each well and incubated for 4 h at 37 °C. Then 150 µL DMSO was added in each well to dissolve the MTT formazan crystals, determined the relative cell viability [10].

### 2.3 Real-time fluorescence qPCR method

Total RNA was extracted from mice lung tissues or A549 cells after Bap treated. The mRNA expression of *HSP90, HSC70* and *Lamp-2a* was detected by qPCR. The method was performed as previously described [11], and the primer listed in Table 1.

**Table 1.**
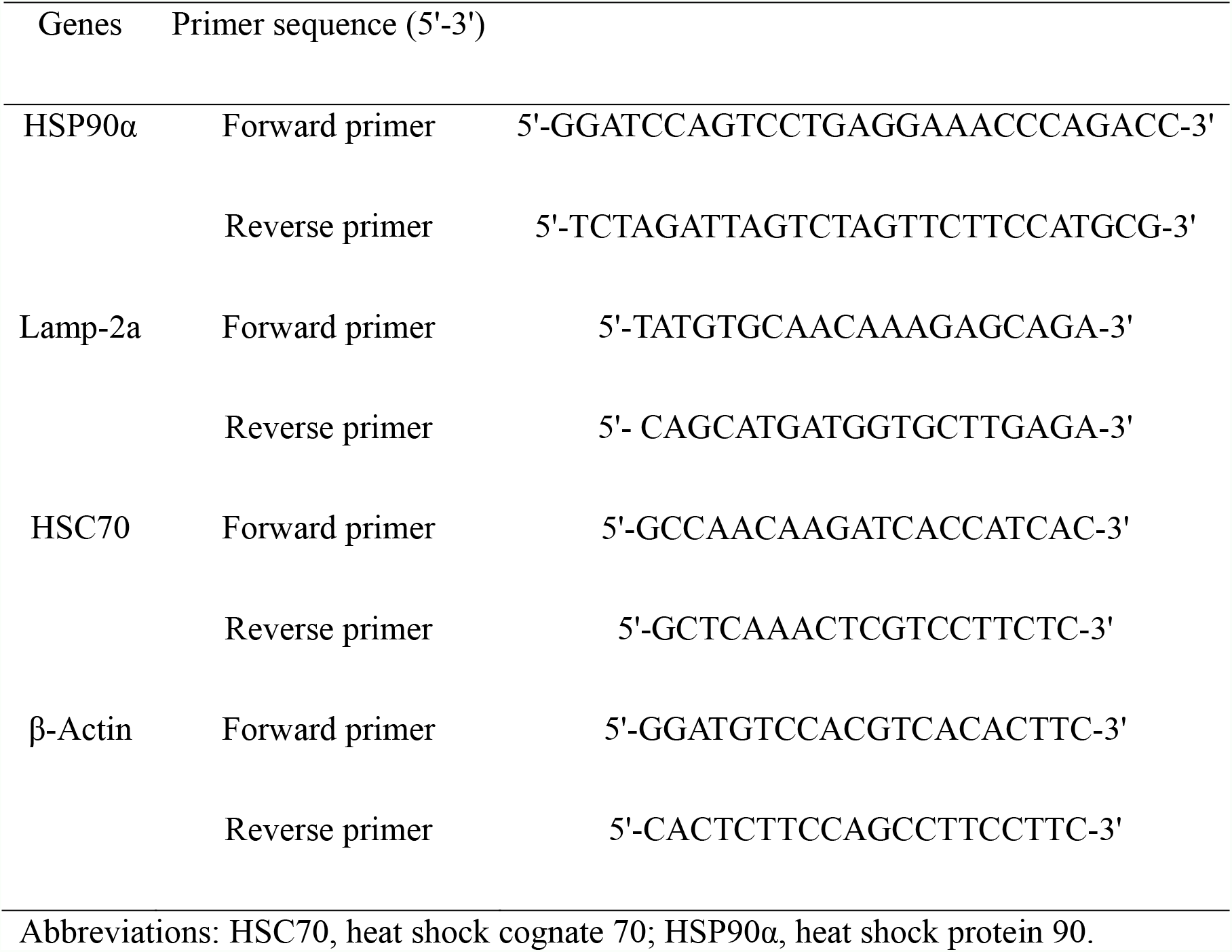
Primer sequences.

### 2.4 Immunohistochemistry

Immunohistochemistry was performed as previously described [12]. Mice lung tissues were prepared according to the instructions of the UltraSensitiveTM S-P hypersensitive kit (MXB, Fuzhou, China). The protein expression of HSP90, HSC70 and Lamp-2a (Abeam, Cambridge, UK) were evaluated double-blind. The staining results were analyzed by the degree of color development and the positive expression rate.

### 2.5 Western blotting

After treatment, A549 cells were lysed, and the protein lysates were subjected to western blotting as described above [13]. The membranes were blocked in 5% non-fat milk in tris-buffered saline buffer, then incubated overnight at 4°C with primary antibodies anti-HSP90, HSC70, Lamp-2a, and GAPDH (Abcam, Cambridge, UK). The second antibody was added, and the membranes were incubated for 1 h at room temperature in the dark. Bound antibodies were analyzed with ImageJ software.

### 2.6 Comet Assay

Single cell gel electrophoresis was performed to determine the effect of Bap on DNA damage.

The protocol was performed as previously described [14]. The slides were stained with ethidium bromide (10 μg/mL) and observed under fluorescent microscope.

### 2.7 γ-H2AX Staining

Monoclonal antibody mouse anti-phospho-H2AX (Abcam, UK, ab26350) was used to detect γ-H2AX foci [15]. Nuclei were stained with 40,6-diamidino-2-phenylindole dihydrochloride. An inverted fluorescence microscope was used to acquire the images.

### 2.8. Construction of HSP90α shRNA plasmid

shRNA target sequences (HSP90α shRNA forward 5’-CACCGTGACTGGGAAGATCACTTGTTCAAGAGACAAGTGATCTTCCCAGTCACTTT TTTG-3’, reverse5’-CGAAGCTTCAAAAAAGTGACTGGGAAGATCACTTGTCTCTTG AACAAGTGATCTTCCCAGTCAC-3’) were designed by Primer 5.0 and ligated into pSGU6/GFP/Neo plasmid as described [16]. In brief, 2 × 10^5^ A549 cells were inoculated in 6-well plates and then transfected using 10 mL Lipofectamine®3000 transfection reagent with 4 mg of the indicated shRNA.

### 2.9 Statistics

The data were expressed as the 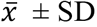, and the Student’s *t*-test was processed by GraphPad Prism software. One-way ANOVA Spearman or non-parametric LSD-*t* analyses were used to distinguish differences between groups. *P* < 0.05 was considered statistically significant.

## 3. Results

### 3.1 The CMA-related genes HSP90α, HSC70 and Lamp-2a are up-regulated in C57BL mice lung tissues and A549 cells after Bap treatment

Compared with the control group, HSP90α, HSC70 and Lamp-2a mRNA expressions of lung tissues in the experimental group were higher by qPCR assay (*P* < 0.05). Protein levels of HSP90α, HSC70 and Lamp-2a in the experimental group have the same significantly altered by immunohistochemical method (*P* < 0.05). Details are as in Fig. S1and Table 2. The results indicating that Bap colud up-regulate HSP90, HSC70, and Lamp-2a in C57BL mice lung tissues.

**Table 2.**
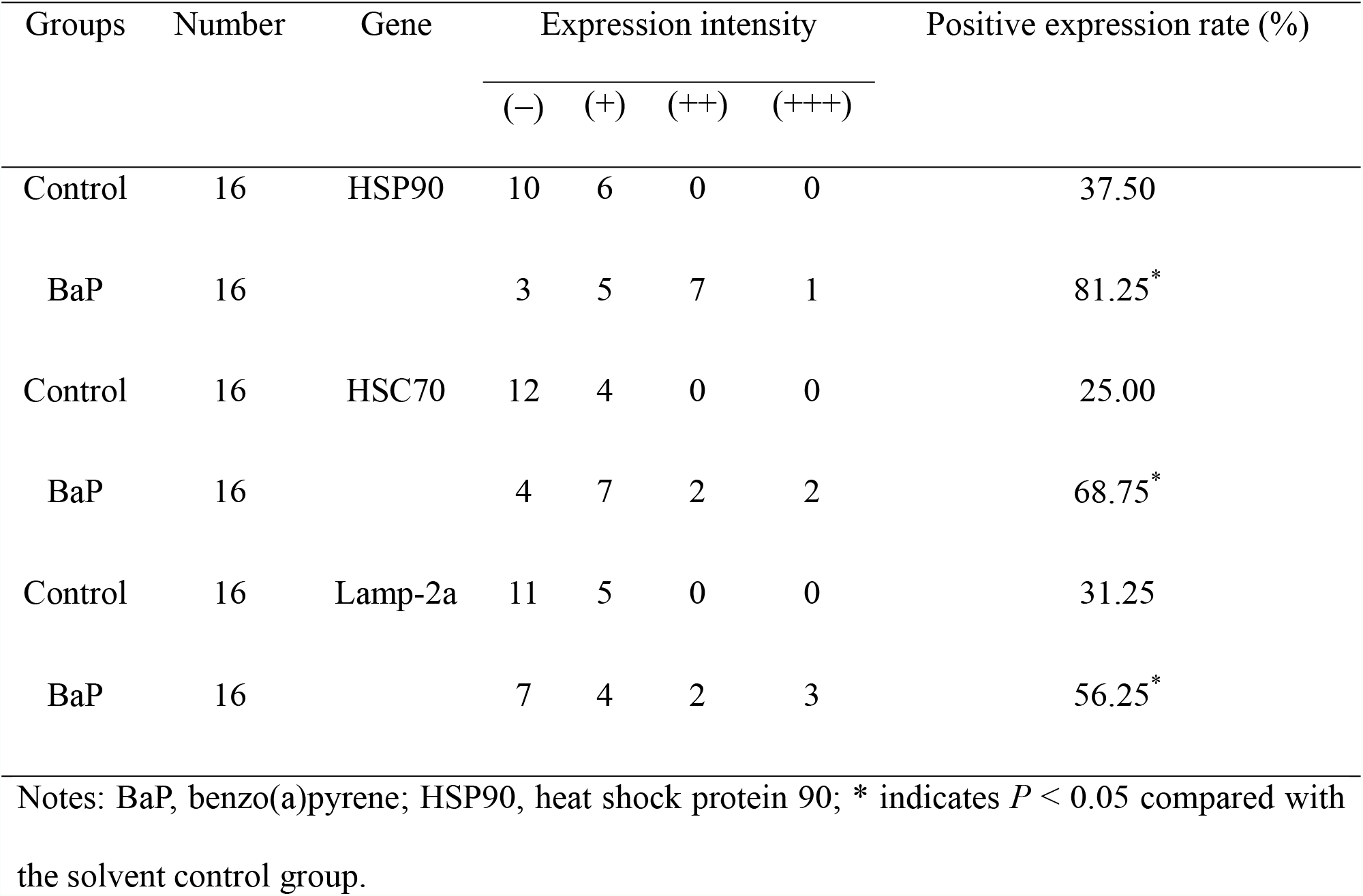
Expression of CMA-associated genes in lung tissues of the control and BaP exposure mouse groups.

| Groups | Number | Gene | Expression intensity |  |  |  | Positive expression rate (%) |
| --- | --- | --- | --- | --- | --- | --- | --- |
|  |  |  | (-) | (+) | (++) | (+++) |  |
| Control | 16 | HSP90 | 10 | 6 | 0 | 0 | 37.50 |
| BaP | 16 |  | 3 | 5 | 7 | 1 | 81.25* |
| Control | 16 | HSC70 | 12 | 4 | 0 | 0 | 25.00 |
| BaP | 16 |  | 4 | 7 | 2 | 2 | 68.75* |
| Control | 16 | Lamp-2a | 11 | 5 | 0 | 0 | 31.25 |
| BaP | 16 |  | 7 | 4 | 2 | 3 | 56.25* |

Notes: BaP, benzo(a)pyrene; HSP90, heat shock protein 90; \* indicates $P < 0.05$ compared with the solvent control group.

### 3.2 Bap-induced CMA is associated with DNA damage induction

We found Bap with 0 to 10μM for 24 h did not affect A549 cells proliferation (Fig. S2A) by MTT.

By alkaline comet assay, the statistics for cell counting results showed that as increasing Bap concentrations, the tailing rate and the average tail length of cells increased significantly (Fig. 1A).

**Fig 1.**
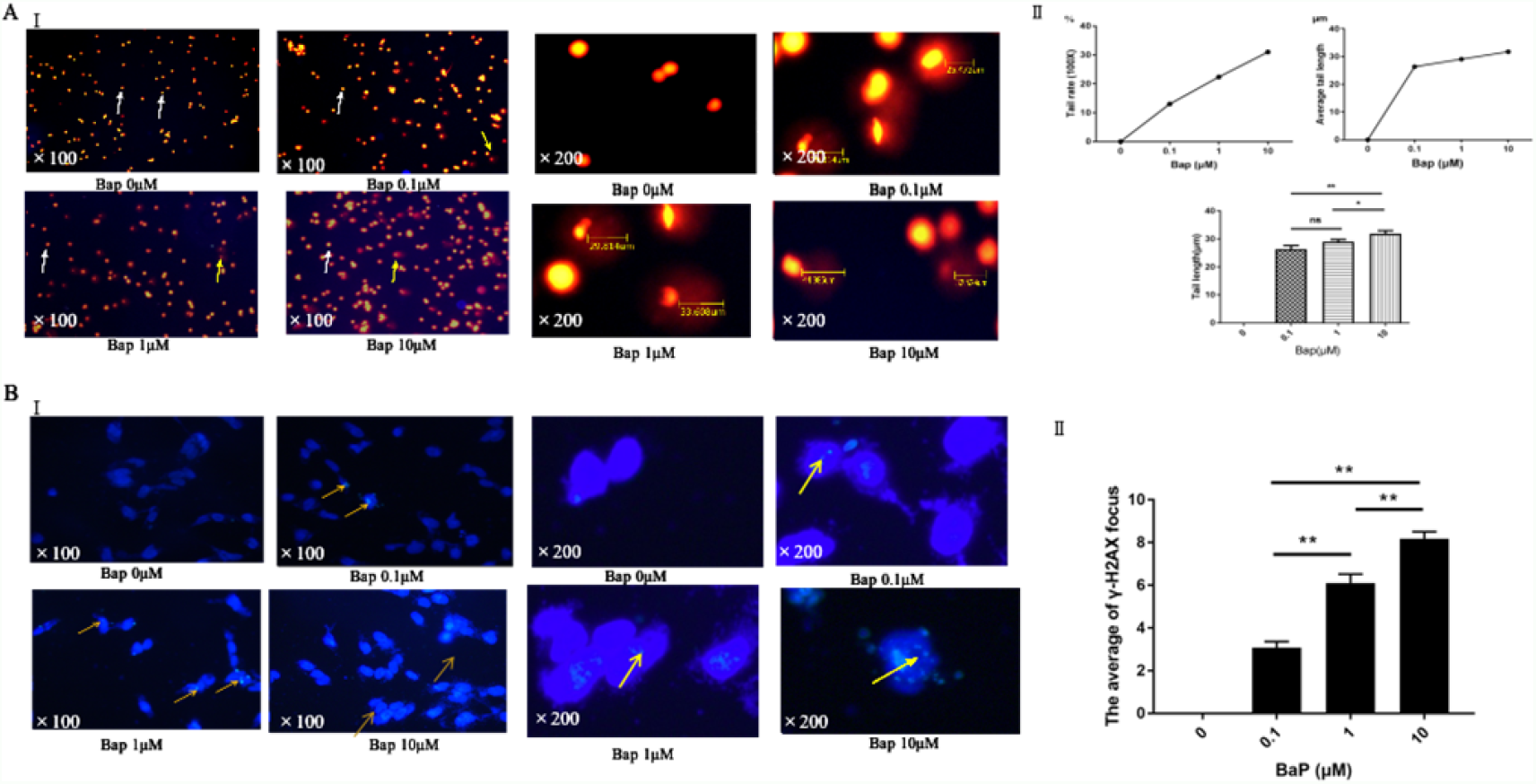
Bap caused DNA damage in A549 cells in a dose-dependent manner. (A) Panel I: The effect of different concentrations which is from 0μM to 10μM Bap on DNA strand breaks in A549 cells was assessed by comet assay; Panel II: Analysis chart of statistical results. (B) Panel I: The effect of 1μM and 10 μM BaP on double-stranded DNA strand breaks in A549 cells was assessed by immunostaining of γ-H2AX foci. Panel II: Analysis chart of statistical results. (\**P* < 0.05; \*\**P* < 0.001).

By immunofluorescence detection of γ-H2AX focus experiment, we found the difference of γ-H2AX focus at the different concentrations have statistically significant (*P* < 0.01). With the increase of Bap concentrations, the mean of γ-H2AX focal points and their average values increased. The results indicated that the degree of DNA double-strand breaks in A549 cells was positive correlation with Bap concentrations (Fig. 1B).

According to qPCR results (Fig. S2B), HSP90, HSC70 and Lamp-2a mRNA expression in A549 cells significantly increased and the differences between groups were statistically significant (*P* < 0.01).

Western blot results showed that HSP90, HSC70 and Lamp-2a protein expressions were significantly increased (Fig. S2C). The difference between groups were statistically significant (*P* < 0.01, *P* < 0.05).

Overall, all these results indicated that Bap could induce CMA-related genes expression (HSP90, HSC70, Lamp-2a) and could promote CMA.

### 3.3. Bap regulates CMA through HSP90

#### 3.3.1 Bap affects CMA-related genes expression by HSP90 inhibitor treated

The results of qPCR and Western blot (Fig. S3A, S3B) showed that compared with the Bap group, expression of HSP90 mRNA and protein remained unchanged in the BaP + NVP-AUY922 group (NVP-AUY922 was a potent HSP90 function inhibitor). And compared with the Bap group, mRNA and protein expressions of HSC70 and Lamp-2a were reduced in the BaP + NVP-AUY922 group, the difference was statistically significant (*P* < 0.05). The results showed that the level of CMA was also inhibited at varying degrees by NVP-AUY922 treated, further illustrated that Bap regulates CMA through HSP90.

#### 3.3.2 Effect of Bap on CMA decreases after gene silencing of HSP90α

The expression levels of HSP90, HSC70, Lamp-2a mRNA and protein in the control group were set to 1. Compared with the shRNA-1 + Bap group, the expression levels of HSP90, HSC70, and Lamp-2a mRNA and protein were not statistically significant in shRNA-1 group (*P* > 0.05). Bap had no effect on the expression of HSP90, HSC70 and Lamp-2a, indicating that Bap regulates CMA through HSP90 (Fig. S3C, S3D).

### 3.4. Bap does not affect DNA damage and no effect on CMA after gene silencing of HSP90α

#### 3.4.1 After gene silencing of HSP90α, to assess the impact of Bap on DNA damage and the effects on HSP90, HSC70, Lamp-2a mRNA and protein expressions we carried out an alkaline comet assay

Under fluorescence microscope, the cell tailing rates of the control, shRNA-1, Bap, and shRNA-1 + Bap groups were 0%, 16.26%, 31.07%, and 19.34%, respectively, with the average smear length of the cells in each group 0μM, 23.46μM, 28.78μM, and 23.95μM, respectively. Compared with the shRNA-1 + Bap group, the tailing rate and average tailing length increased, however, the differences were not statistically significant (Fig. 2A). It was shown that after gene silencing HSP90α, the impact of Bap on DNA damage was weakened.

**Fig 2.**
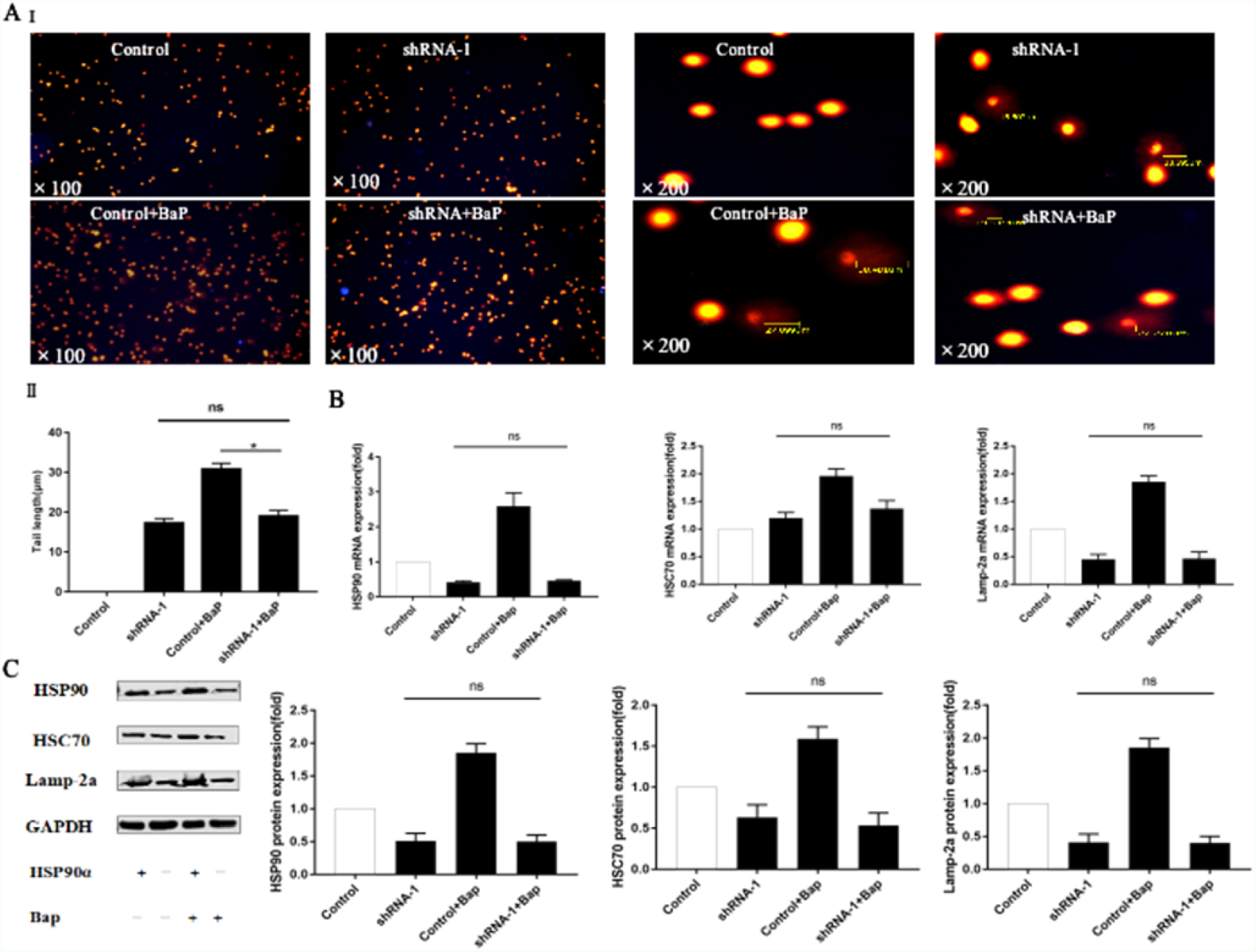
BaP reduces DNA double stranded after HSP90α silencing, and does not affect CMA. A shows BaP treatment silenced HSP90 A549 cells to produce comet tails through the alkaline comet test. I means the tailing rate and average tailing length of A549 cells under microscope; II means analysis chart of statistical results. B and C show mRNA and protein expression of CMA-related genes (HSP90, HSC70 and Lamp-2a) by BaP after HSP90α silencing.(ns means *P* > 0.05, \**P* < 0.05).

Western blot and qPCR analysis for the indicated proteins and mRNA in the cells described above. Compared with Bap group and shRNA-1 group, HSP90, HSC70, Lamp-2a mRNA and protein expressions were decreased in shRNA-1 + Bap group. The differences were not statistically significant (*P* > 0.05). It was shown that Bap could not affect CMA after gene silencing of HSP90α (Fig. 2B, 2C).

#### 3.4.2 After gene silencing of HSP90α, γ-H2AX foci staining was used to detect the degree of DNA double-strand breaks, and the effect on HSP90, HSC70, and Lamp-2a mRNA and protein expression

We classified A549 cells into distinct groups by different treatments: the control, shRNA-1, Bap and shRNA-1 + Bap groups, carried out fluorescence microscope. Compared with the Bap group, the average value of foci of γ-H2AX was significantly reduced in the shRNA-1 + Bap group (*P* < 0.05). Compared with the shRNA-1 group, there was no significant difference in the average number of γ-H2AX focal points in the shRNA-1 + BaP group (*P* > 0.05). The results suggested after *gene silencing of HSP90α*, the damage of Bap to DNA double-strand breaks was reduced (Fig. 3A).

**Fig 3.**
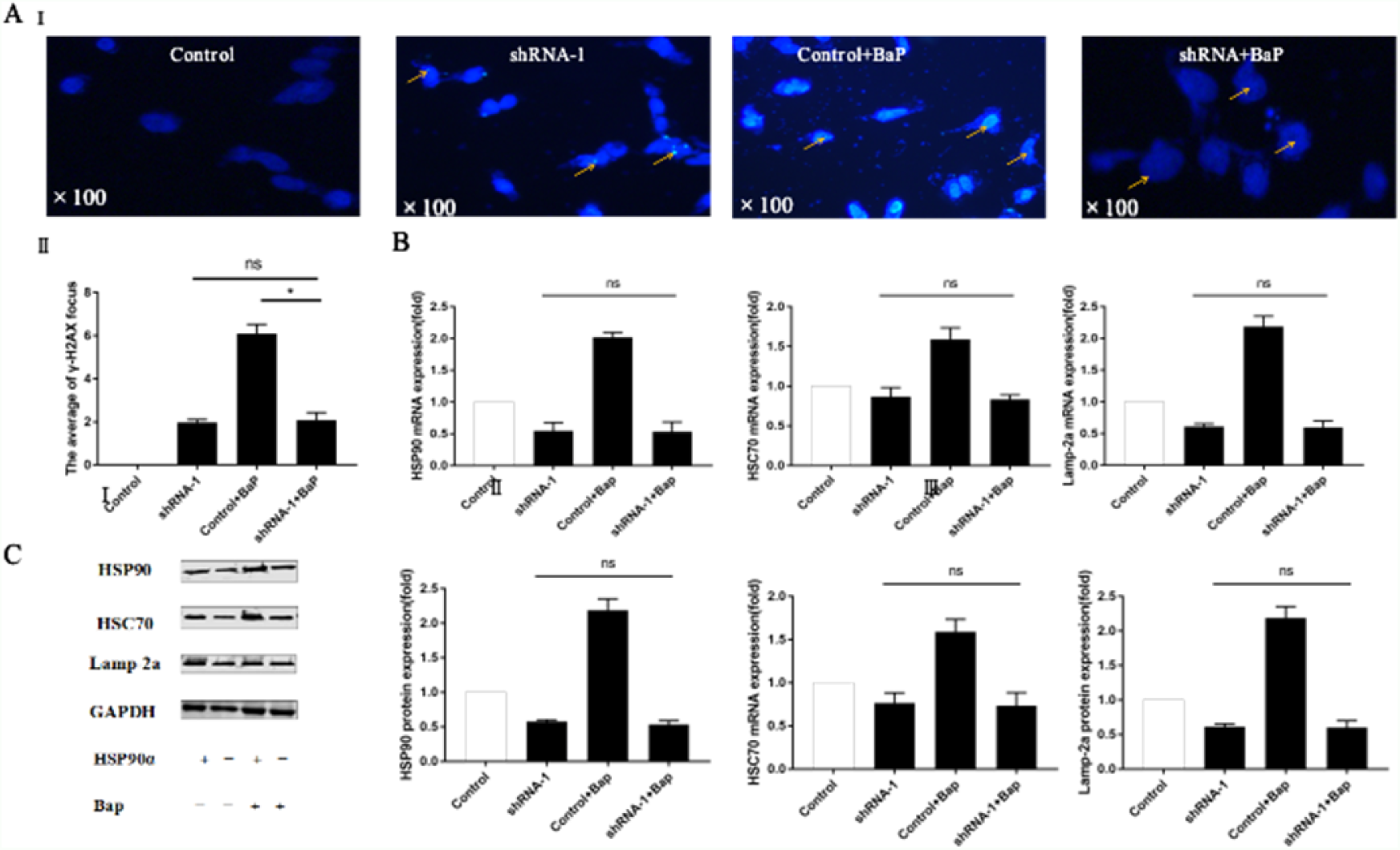
BaP reduces DNA damage after HSP90α silencing, and does not affect CMA. A shows γ-H2AX focus number of silenced HSP90α A549 cells treated with BaP. I means different concentrations of BaP on silenced HSP90 A549 cells under microscope; II means analysis chart of statistical results. B and C shows mRNA and protein expression of CMA-related genes (HSP90, HSC70 and Lamp-2a) by BaP after HSP90α silencing (ns means *P* > 0.05, \**P* < 0.05).

Western blot and qPCR analysis for the indicated proteins and mRNA in the cells described above. Compared with Bap group and shRNA-1 group, HSP90, HSC70, Lamp-2a mRNA and protein expressions were decreased in shRNA-1 + Bap group. The differences were not statistically significant (*P* > 0.05). The results showed that after gene silencing of HSP90α, the damage of Bap to DNA double-strand breaks was reduced and further reduced the effect on CMA (Fig. 3B, 3C).

## 4. Discussion

To explore the effect of Bap on CMA, we conducted Bap gastric gavage in live C57BL mice to assess the CMA-related gene expressions in lung tissues. The results showed that Bap up-regulated the expression of CMA-related genes (*HSP90, HSC70* and *Lamp-2a*), indicating that Bap promotes CMA. It had been reported in the literature that smoking could cause increased expression of autophagy-related genes[17]. Bap was the main component of cigarette smoke, which indirectly supports our research.

According to the results of comet assay and γ-H2AX staining, as the concentration of Bap increase, DNA damage degree were increased. Our results are consistent with other published report[8]. Interestingly, HSP90, HSC70 and Lamp-2a mRNA and protein also increased to a similar extent in the same cells. The results indicated that Bap could cause DNA damage and affect CMA-related genes, further demonstrated that Bap could promote CMA.

As a form of selective autophagy, CMA is rarely studied, and its molecular mechanism is unknown [18]. HSP90 is an important molecular chaperone and can mediate CMA[19]. However, what role it plays in the process of Bap-induced CMA has not been reported. On the basis of our above results, our propose the possibility of Bap regulates CMA through HSP90. To validate this conjecture, we conducted several experiments and observations supported this conjecture. Firstly, compared with the Bap group, mRNA and protein expressions of HSC70 and Lamp-2a were reduced in the HSP90 inhibitors (NVP-AUY922) + Bap group, the difference was statistically significant (*P* < 0.05). (compared with the control group?) It is reported from the literature that NVP-AUY922 has a better inhibitory effect on HSP90 than other inhibitors [20]. Since NVP-AUY922 can only inhibit the function of HSP90 by binding to the ATP pocket of HSP90, it cannot inhibit the expression of HSP90 protein and mRNA [21]. In addition to HSP90 acting as a molecular chaperone in cells, HSP90 can also be secreted into cells, and secreted HSP90 has been confirmed to be HSP90α [22]. Next, we used RNA interference technology to silence HSP90α to observe the effect of HSP90 on CMA. The results showed that compared with the shRNA-1 + Bap group, the expression levels of HSP90, HSC70, and Lamp-2a mRNA and protein were not statistically significant in shRNA-1 group (*P* > 0.05). In addition, we found that silencing HSP90α reduced HSP90 protein expression by approximately 55%, indicating that HSP90α accounted for the majority of HSP90 isoforms in A549 cells. Thus, the results show, after HSP90 is inhibited or HSP90α is silenced, Bap-induced CMA is also inhibited to varying degrees, indicating that HSP90 plays an important role in Bap-regulated chaperone autophagy. Further, we took 1mL of previously remaining cell suspension in each group, which were detected mRNA and protein expression of CMA-related genes by qPCR and Western blot respectively. Simultaneously the DNA damage degree in each group were detected by the γ-H2AX focus. Ultimately, there were no significant difference of DNA damage degree, the mRNA and protein expression of CMA-related genes in shRNA-1 +Bap group and the shRNA-1 group. Collectively, our results suggest that Bap induce DNA damage, which promotes CMA.

From the results of this research, Bap regulated CMA through HSP90 thought enhanced HSP90, and Bap induce DNA damage, which promotes CMA. Ernst *et al*.[23] showed that inhibition of HSP90 significantly affected DNA repair and they confirmed that HSP90 had an pivotal role in DNA damage. Possibly due to multiple components involved in DNA double-strand break repair mechanisms (including BRCA1, BRCA2, CHK1, DNA-PKcs, FANCA, and MRE11 / RAD50 / NBN complexes) were described as HSP90 client proteins, and they are all DNA damage Regulators [24-25]. Robert *et al*.[26] showed that autophagy could be activated by DNA damage, and DNA damage could affect autophagy. Previous researches mostly focused on the relationship between HSP90 and autophagy, as well as autophagy with DNA damage [27-28]. However, the relationship between HSP90, CMA, and DNA damage is still elusive. Our research confirmed that Bap promoted CMA by HSP90, with the process that HSP90 participated DNA damage caused by Bap. The study provides a clearly evidence that Bap regulates CMA via HSP90 and a sight of the mechanism of Bap action.

## Acknowledgments

Not applicable.

## Abbreviations

BaP: benzo(a)pyrene;
CMA: chaperone-mediated autophagy;
HSC70: heat shock cognate 70;
HSP90: heat shock protein 90;
Lamp-2a: lysosomal-associated membrane protein type 2 receptor;
MTT: tetramethylazozolium salt;
q-PCR: quantitative PCR;
shRNA: short hairpin RNA

## Supplementary Figure Legends

**Fig. S1.**
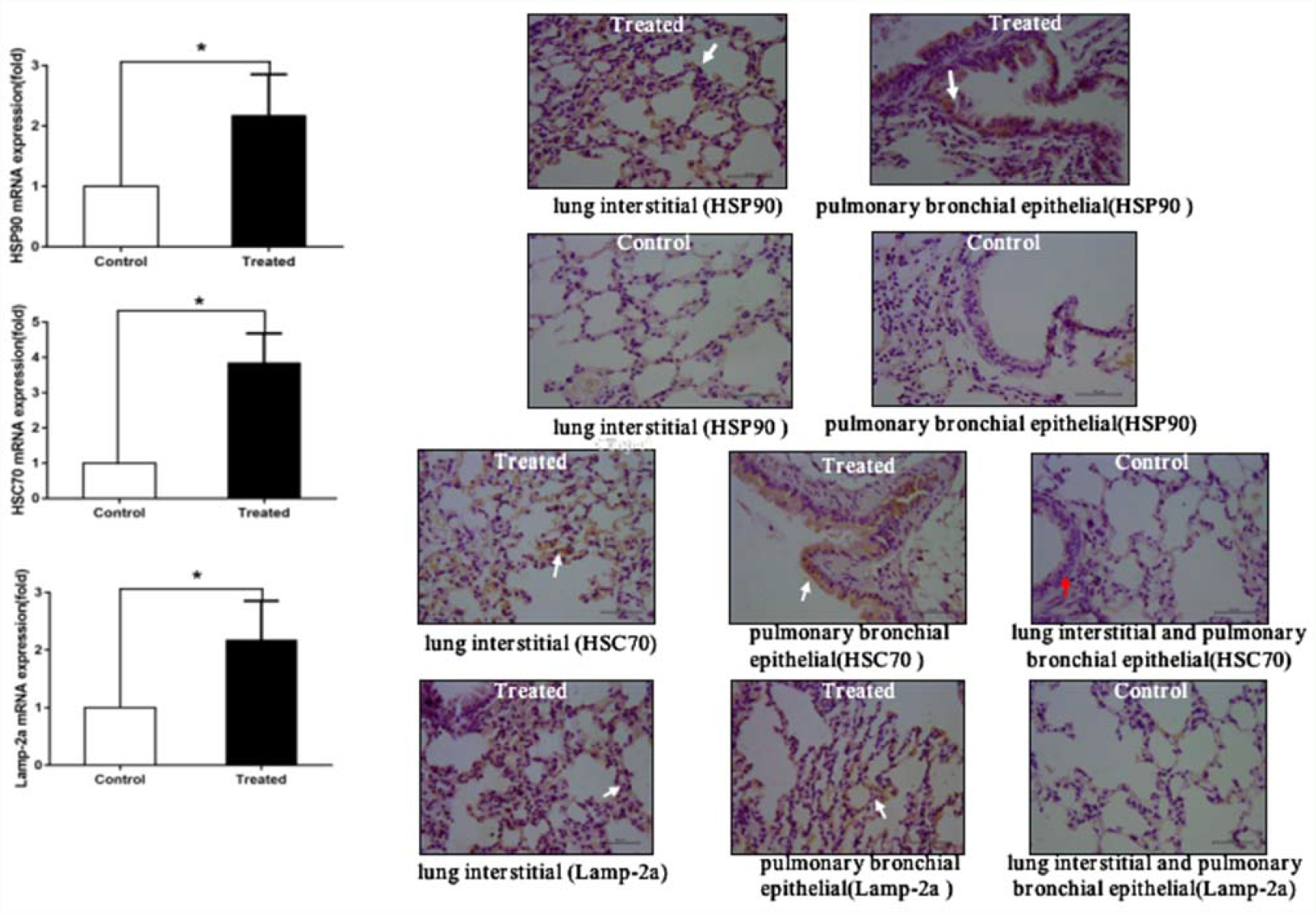
Effect of BaP on the expression of CMA-related genes in lung tissue of C57BL mice. (A) mRNA expression of HSP90, HSC70 and Lamp-2a in lung tissues of control C57BL mice and mice that were treated with BaP (\**P*<0.05). (B) Representative results from HSP90, HSC70 and Lamp-2a immunohistochemistry staining. White arrows indicate the cytoplasmic portion of the cell.

**Fig. S2.**
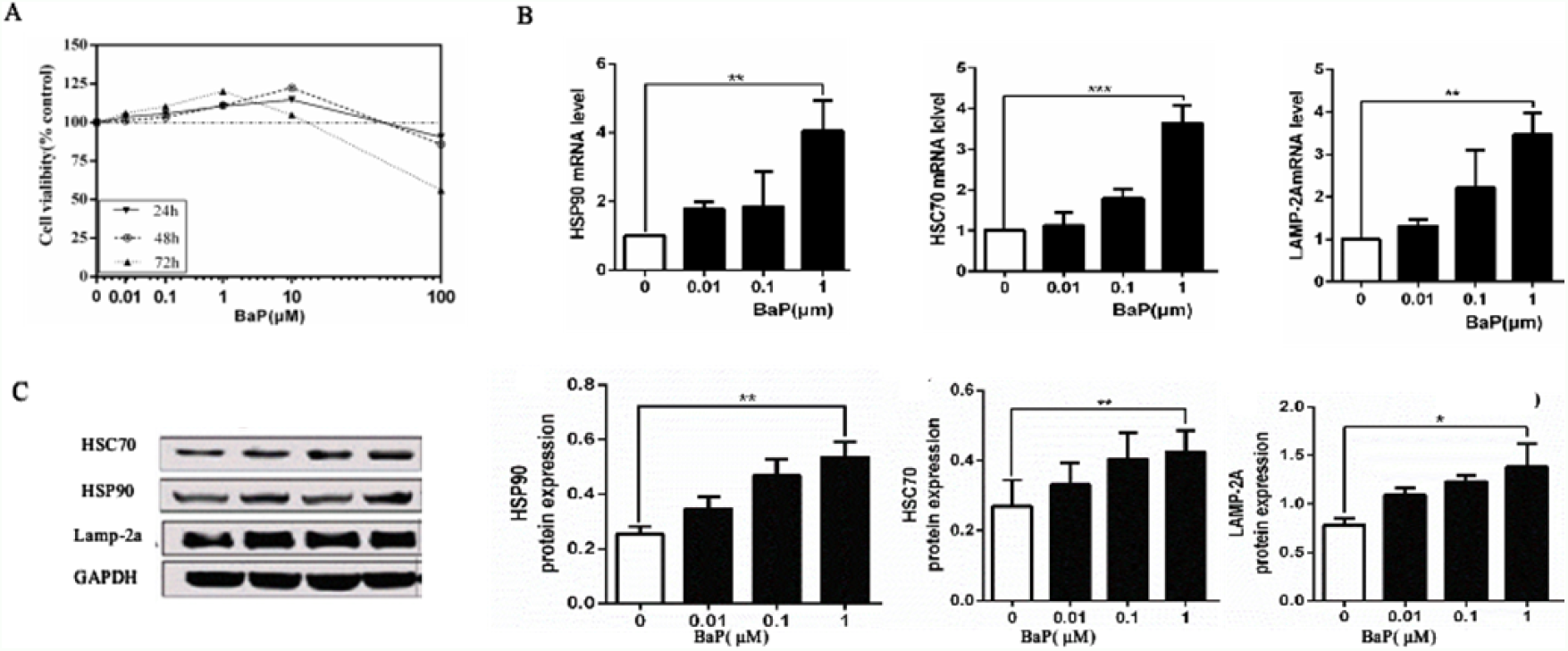
BaP promotes expression of CMA-related genes. A and B show the effects of different concentrations of BaP on the activity of A549 cells at different time.

**Fig. S3.**
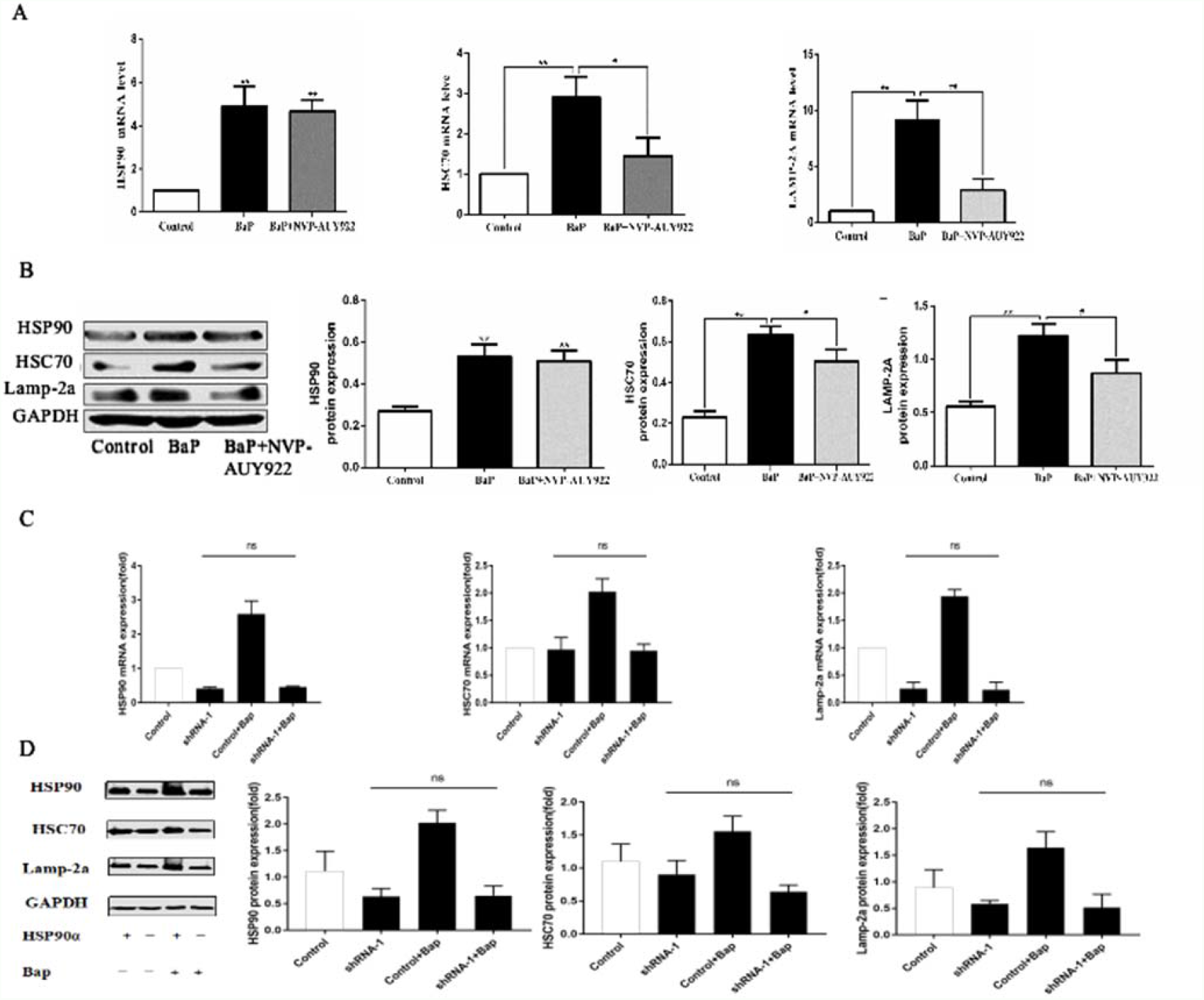
BaP regulates CMA through HSP90. A and B show the mRNA and protein expression levels of HSP90, HSC70 and Lamp-2a in A549 cells treated with NVP-AUY922 and BaP. C and D show the mRNA and protein expression levels of HSP90, HSC70 and Lamp-2a in silent HSP90α A549 cells treated with BaP (ns means *P* > 0.05, \**P* < 0.05, \*\**P* < 0.001).

